# Greater spear nosed bats commute long distances alone, rest together, but forage apart

**DOI:** 10.1101/2021.09.30.462631

**Authors:** M. Teague O’Mara, Dina K.N. Dechmann

**Affiliations:** Bat Conservation International; Department of Biological Sciences, Southeastern Louisiana University; Department of Migration, Max Planck Institute of Animal Behavior; Smithsonian Tropical Research Institute; Department of Biology, University of Konstanz; Centre for the Advanced Study of Collective Behaviour, University of Konstanz

**Keywords:** GPS tracking, movement ecology, energy expenditure, foraging, social behaviour

## Abstract

Animals frequently forage in groups on ephemeral resources to profit from social information and increased efficiency. Greater spear-nosed bats (*Phyllostomus hastatus*) develop group-specific social calls, which are hypothesized to coordinate social foraging to feed on patchily-distributed balsa flowers. To test this, we tagged all members of three social groups of *P. hastatus* on Isla Colón, Panamá, using high frequency GPS during a season when balsa had begun to flower. We found commuting distances of 20-30 km to foraging sites, more than double of what has been previously reported. In contrast to our expectations, we found that tagged individuals did not commute together, but did join group members in small foraging patches with high densities of flowering balsas on the mainland. We hypothesized that close proximity to group members would increase foraging efficiency if social foraging were used to find flower clusters, but distance between tagged individuals did not predict foraging efficiency or energy expenditure. However, decreased distance among tagged bat positively influenced time outside of the cave and increased the duration and synchrony of resting. These results suggest that social proximity seemed to be more important during resting, and indicate that factors other than increased feeding efficiency may structure social relationships of group members while foraging. It appears that, depending on the local resource landscape, these bats have an excellent map even of distant resources and may use social information only for current patch discovery. They then may no longer rely on social information during daily foraging.

**Highlights:**

- Unlike in Trinidad, greater spear-nosed bats in Panamá do not commute or forage as a group
- Foraging distances are double of previously known and across the sea
- Bats rest near each other, but do not forage more efficiently when close
- Social information may mainly be used for the discovery of feeding areas
- Local resource landscapes may cause strong variation in social behaviour

## Introduction

Animals must respond to changes in the spatiotemporal distribution of resources, and their movement decisions to search for and exploit food resources directly impact the ability to satisfy dietary requirements and affects their fitness (Bell, 1990). When resources are ephemeral (e.g., spatially patchy, temporally unpredictable), using social information while moving with others may help them find resources more efficiently (Bhattacharya & Vicsek, 2014). For example, fish track moving refugia by matching speed to group mates (Berdahl, Torney, Ioannou, Faria, & Couzin, 2013), insect- and fish-eating bats converge on the feeding calls of conspecifics (Dechmann et al., 2009; Egert-Berg et al., 2018), seabirds follow the white plumage of foraging flocks (Beauchamp & Heeb, 2001), and penguins are able to capture more fish when foraging together (McInnes, McGeorge, Ginsberg, Pichegru, & Pistorius, 2017).

Foraging in groups can convey energetic benefits by increasing foraging success and make energy intake more reliable (Giraldeau & Beauchamp, 1999; McInnes et al., 2017; Snijders et al., 2021). Individuals are required to maintain cohesion and spatiotemporal coordination to benefit from interactions with conspecifics (Conradt & Roper, 2005). While maintaining strong social bonds can provide long-term fitness benefits (Bohn, Moss, & Wilkinson, 2009; Silk et al., 2010), moving with group members can increase feeding competition and the immediate costs of transport (Usherwood, Stavrou, Lowe, Roskilly, & Wilson, 2011). Less time and energy spent on finding food patches due to information provided by group members may be especially important for species foraging on ephemeral resources, but little evidence is available. The use of social information should decrease energy expenditure and / or increase foraging efficiency and result in higher rates of return during foraging bouts.

Bats are an excellent group to test how resource ephemerality and energy expenditure direct group foraging. They spend large proportions of their energy budget on locomotion that is fueled by the food of the day (O’Mara et al., 2017), and feed on resources that are often widely dispersed and unpredictable. Bat species that forage for ephemeral insect swarms eavesdrop on the echolocation buzzes that reveal a capture attempt (Dechmann et al., 2009; Dechmann, Kranstauber, Gibbs, & Wikelski, 2010; Egert-Berg et al., 2018). Beyond this often-opportunistic behaviour, some species will also search food near group members to maximize the discovery of feeding patches (Dechmann et al., 2009; Dechmann et al., 2010; Egert-Berg et al., 2018). Bats of many species readily incorporate social information about food across a range of cues (O’Mara, Dechmann, & Page, 2014; Page & Ryan, 2006; Ramakers, Dechmann, Page, & O’Mara, 2016; Ratcliffe & ter Hofstede, 2005; Wright, 2016), and the nature of the resource they feed on as well as how tightly they depend on it can be used to predict if and when social information should be used during foraging (Kohles, O’Mara, & Dechmann, 2022).

Greater spear-nosed bats (*Phyllostomus hastatus*) seasonally feed on an ephemeral resource, the nectar of balsa trees (*Ochroma pyramidale*). In a well-studied population on Trinidad, female *P. hastatus* form stable life-long groups of unrelated females (McCracken & Bradbury, 1981; Wilkinson, Carter, Bohn, & Adams, 2016). They synchronize reproduction, converge on a loud group-specific call that requires extended learning, and perform several cooperative behaviours at the group level, such as babysitting and pup guarding from the infanticide attempts of neighbouring groups (Boughman & Wilkinson, 1998; Boughman, 1998; Wilkinson & Boughman, 1998; Wilkinson et al., 2016). They are omnivorous, but during the dry season, they feed nearly exclusively on balsa nectar. These pioneer trees are a rare and patchy resource and the group-specific social calls are hypothesized to recruit group members to flowering trees to exploit or defend them collectively (Wilkinson & Boughman, 1998). However, the number of flowers available on a given tree is limited (Kays et al., 2012), many other animals feed on them, and the energy requirements of these bats are large (Kunz, Robson, & Nagy, 1998). Thus, the potential reasons for recruiting others to these flowers warrant further investigation, especially if *P. hastatus* social groups forage together to feed on these flowers. In addition, the availability of balsa and thus the value of social information (Kohles et al., 2022) may vary locally, and it is unclear if group foraging and resource defence occur across the species’ range.

We used high frequency GPS loggers to tag three groups of *P. hastatus* in Panamá and recorded complete foraging trips. We used these GPS data to construct proximity-based social networks to test how social associations are linked with foraging performance and behaviour. We hypothesized that, like in Trinidad, *P. hastatus* forage in groups during the dry season. We thus expected them to commute to a food source socially (either as a group or subgroups of individuals) and exploit it together. We also hypothesized that social foraging increases foraging success, and that proximity that is within hearing distance of social calls increases foraging efficiency and lower energy expenditure despite potential competition trade-offs. With this study we make an important contribution to how foraging behaviour may vary across sites and seasons, and thus the intricate link between a local resource landscape and the resulting social behaviour.

## Methods

Data were derived from 39 adult *Phyllostomus hastatus* (37 F / 2 M) that were captured from three roosting groups in a cave (“La Gruta”) on Isla Colón, Bocas del Toro, Panamá using a bucket trap. Groups were captured sequentially and there was no overlap among groups in the nights they were tracked. Roosting groups were individuals co-roosting in small depressions on the cave ceiling, consistent with previous work in Trinidad. All bats from these groups were fitted with GPS tags, with the exception of two females that had heavily worn teeth and old, severe injuries. Both males were adult harem males, the females comprised 15 nulliparous young females, and 19 postlactating females. GPS tag retrieval success varied among groups (Group 1: 13 F / 1 M deployed with 5 F / 1 M or 42.8% recovered; Group 2: 6 F deployed with 4 F or 66.7% recovered; Group 3: 16 F / 1 M deployed with 8 F / 1M or 52.9% recovered). Bats from a single social group were placed into a wire mesh cage covered with a breathable cotton cloth where they roosted together calmly until removed for processing. Bat mass was recorded to the nearest 0.5 g, forearm length measured to the nearest 0.1 mm, and each bat received a subcutaneous PIT tag (ID100; Euro ID, Frechen, Germany). To measure wing dimensions for flight power estimates, we took photos of one fully outstretched wing placed flat over mm graph paper. Bats were fitted with a GPS data logger (Gypsy-5 GPS, TechnoSmart, Rome, Italy (O’Mara et al., 2021)) that was wrapped in clear shrink tube. The logger was mounted on a silk collar (0.8 cm wide) and closed with Safil-C degradable suture (Aesculap/B. Braun, Co, Tuttlingen, Germany, (O’Mara, Wikelski, & Dechmann, 2014)). Total collar + GPS weight was 6.8 ± 0.51 g, which represented 5.7 ± 0.4 % of body mass (range: 4.5 - 6.6 %).

GPS tags were programmed to collect location fixes from 18 h – 06 h local time every one or two s. When there was not adequate GPS reception, tags went into a low energy sleep state for five mins and then restarted to search for satellites for 90 s. Tag function varied due to the deep cave roost used by the bats and resting under presumably dense foliage while foraging. We retrieved 18 tags with analyzable data: five females and one male from group one, three females from group two, and eight females and one male from group three (Table S1). The 18 recovered tags collected one to four nights of data for a total of 34 bat nights. We removed from analysis five nights from five different bats where fewer than 30 mins were tracked for various reasons (e.g., the bat remained in the cave for most of the night draining the battery), leaving 29 bat nights from 16 females and 2 males with a range of 75.5 – 307.5 min of data collected per night (mean ± sd: 197.31 ± 60.35, Table S1). We tagged entire social groups and retrieved 42-67% of the tags per group. Inferences made then reflect only social interactions reconstructed from these retrieved tags and not the remainder of the social group, all other social groups in the cave, or all other caves from the surrounding area.

Decisions to forage with group members may rely on the overall energetic costs relative to feeding success of social foraging. We use airspeed to estimate instantaneous energy expenditure based on well-established flight power estimates (Pennycuick, 2008) To estimate flight airspeed and subsequent energy expenditure, wind data were collected at an automated weather station (9.351, −82.258) at 15-min intervals by the Physical Monitoring Program at the Smithsonian Tropical Research Institute for their Bocas del Toro field station and downloaded from http://biogeodb.stri.si.edu/physical_monitoring/research/bocas. Wind speed and direction were collected every 10 s with a RM Young Wind Monitor Model 05103. Mean wind speed and wind direction were then calculated at the end of every 15-min interval.

## Analysis

All analyses were conducted in R 4.2.1 (R Core Team, 2022).

### Ground speed, wind speed, and wind accommodation

Ground speed (speed of movement relative to the ground) and bearing were calculated for successive time points in the *move* package (Kranstauber, Smolla, & Scharf, 2018). To calculate airspeed (speed of movement relative to the moving air column), wind speed and direction were annotated for each GPS location using a weighted interpolation of the U and V components of the available 15 minute wind samples to match the higher resolution (0.5 – 1 Hz) of the GPS sampling (O’Mara et al., 2021; O’Mara et al., 2019; Safi et al., 2013). Wind support was calculated as the length of the wind vector in the direction of the bat’s flight where positive values represent tailwind and negative values headwind and are given as total support in m s^-1^. Crosswind was calculated as the absolute value of the speed of the wind vector perpendicular to the travel direction, and airspeed was calculated as the square root of [(ground speed – wind support)^2^ + crosswind^2^].

### Behavioural segmentation

To identify behavioural states of resting, slow foraging flight (i.e., feeding), moving between patches, and commuting, we applied a four-state hidden Markov model in *momentuHMM* (McClintock, Michelot, & Goslee, 2018). These behaviours were chosen as they represent the most likely identifiable biologically-meaningful behaviours derived from time series of GPS locations (de Weerd et al., 2015; Edelhoff, Signer, & Balkenhol, 2016; Gurarie, Andrews, & Laidre, 2009; Gurarie et al., 2015). While the statistical identification of behavioural states is executed through a hidden Markov model (or machine learning or neural network), assignment to a particular behaviour of interest relies on recognizing the limitation of GPS data (time and location) and knowledge of how species move through an environment. Our four behavioural categories differed as follows: bats at “rest” (i.e., no movement) accumulated substantial GPS error and there appeared to be erratic ‘movement’ in the data, with high step lengths (distances between GPS locations) at high turning angles (de Weerd et al., 2015). Slow “foraging flight” had short step lengths with high turning angles, reflecting a slow searching flight pattern with short pauses. “Move” was more directed flight between patches, with increasing step lengths and lower turning angles, and “commute” included highly directed flight where bats were moving quickly with little deviation in their flight paths.

Behavioural states were entered into the hidden Markov model in order of increasing speed and decreasing turning angular mean (i.e., slow flight had larger turning angles, commuting flight was fast with high concentrated turning angles near zero), with step lengths modelled with a gamma error distribution and turning angles with a wrapped Cauchy distribution. Models were fit for each bat on each bat night, and each resulting model was visually inspected to ensure reasonable classification. The track for each bat night was first regularized to one-second intervals using a correlated random walk procedure in function *momentuHMM::crawlWrap* and then passed to the hidden Markov model. We used simulations to target the number of identifiable states and identify the starting values for each state (McClintock et al., 2018; Michelot et al., 2017), and these simulations showed that four-state model always performed better (had lower AIC values) than three- or two-state models. On occasion, a five-state model had better fit although it was often difficult to discern biological meaning between the additional state that was placed very close to the low speed and higher turning angle behaviours of slow foraging flight.

We used a patch approach to identify foraging and resting (i.e., night roost) areas since aggregations of GPS locations should indicate a site of behavioural interest. For each bat night a patch was defined as a cluster of GPS locations that were classified as foraging (slow flight or moving) or as rest. These clusters were identified using density-based spatial clustering of applications with noise (DBSCAN) using function *fpc::dbscan* (Hennig, 2020; Schubert, Sander, Ester, Kriegel, & Xu, 2017) with a minimum of 15 points per cluster at a maximum spatial distance (eps) of 10 m among nearest neighbours. This distance was chosen through visual inspection of diagnostic plots in function *dbscan*::*kNNdistplot* (Hahsler, Piekenbrock, & Doran, 2019). To facilitate spatial comparisons across all individuals in the sequentially tracked groups, we labelled patches with centroids that were less than 30 m apart as a single patch regardless of the night on which they were used. This distance was chosen as there was a clear break in the distribution of pairwise distances among patch centroids and 30 m is slightly larger than the approximate diameter of a balsa crown. These patches were also ground-truthed to evaluate potential plant food composition and presence.

Once foraging patches were identified, we then further classified feeding locations that were likely flower clusters. We used the same DBSCAN procedure on the GPS locations within each foraging patch per bat night, to identify a flower cluster as a position with a minimum of six points per cluster within a maximum spatial distance of 0.8 m. This distance was chosen based on the spatial distribution of flowers observed within *O. pyramidale* (personal observation). We used these likely feeding clusters to define foraging efficiency as feeding clusters divided by the total time tracked (in mins) per night.

### Energy expenditure

The speed that an animal flies in an air column (airspeed) is the most important predictor of its mechanical power output and subsequent total metabolic power output. We estimated the mechanical power of flight in Watts (P_mech_) following Pennycuick (2008) using calculated airspeeds, the capture mass of the animals, and wing length taken from each bat’s wing photo multiplied by two. An individual’s average power curve was generated for each bat at the mean flight altitude (50 m) and 25°C (Figure S1) and returned estimates across all individuals for the minimum power speed of 6.81 ± 0.21 ms^-1^ and maximum range speed of 11.0 ± 0.35 ms^-1^. Minimum power speed represents the most efficient instantaneous flight speed and maximum range speed maximizes the range covered over ground per unit energy expended. In general, bats should fly at their minimum power speed when moving short distances and at the maximum range speed when moving long distances (Hedenström, 2003). The airspeeds used by bats (7 – 9 m s^-1^) falling well within these estimates for energy-efficient flight. To estimate energy expenditure for each night, the instantaneous power output during flight was calculated for each GPS location at the observed altitude and airspeed. Mechanical power output alone underestimates metabolic power requirements (Pennycuick, 2008; von Busse, Swartz, & Voigt, 2013; Ward et al., 2001). To estimate total metabolic power required, we estimated the metabolic power of flight (P_met_) following Ward et al (2001) using the mean estimated flight muscle partial efficiency (E_FM_) for *P. hastatus* in a wind tunnel (0.24667, range: 0.13 – 0.34, (Thomas, 1975)). Total metabolic power was then calculated as: P_met=_ 1.1[(P_mech_ / E_FM_) + P_BMR_]. For locations where the bat was at rest (including while in the cave), we substituted the resting metabolic rate (P_BMR_) of 23.8 J g^-1^ h^-1^ (McNab, 1969) and converted this to 0.0661 W g^-1^. These values were summed to daily energy expenditure (DEE) and compared to DEE values from smaller *P. hastatus* in Trinidad measured through doubly labelled water (Kunz et al., 1998).

To estimate energy returns from foraging, we used the energy content per ml of balsa nectar and the likely flower density per tree (Kays et al., 2012). Total nectar produced by a flower is estimated at 25.5 ml, and balsa nectar sugar concentration decreases over the night from 13.3% at 18 h to 7.9% at 06 h, with an average concentration of 12.4% total sugars (Kays et al., 2012). This is 0.124 g sucrose ml^-1^ nectar that yields 0.47988 kcal ml^-1^ (3.87 kcal g^-1^ sugar *0.124 g ml^-1^). Balsa nectar then has an energy density of 2.007818 kJ ml^-1^. Flowers open with 4.9 ± 1.3 ml of nectar, and there is a sharp decline in nectar production over the night. We assumed that bats drink the full nectar volume present when flowers open (5 ml), which likely over-estimates the amount of nectar truly ingested during feeding events. Peak flower availability is approximately 60 flowers per patch or 5 flowers per m^2^. However, the mean is 20 flowers in a patch and 2.5 flowers per m^2^, with flower density following a normal distribution over the season (Kays et al., 2012).

### Social Proximity Effects on Behaviour

To test the effects of distance to a nearest neighbour on behaviour we used the distance among individuals as a dynamic metric that could change with every second of tracking. We excluded commuting behaviour from this analysis after inspection of the data showed that individuals did not commute together with group mates. We used this pairwise distance to further identify if the changing proximity between individuals affects movement decisions. We limited the potential distance that behaviour could be affected by another individual to 290 m, which is the potential perceptual distance of *P. hastatus* social calls at a peak call frequency of 6725 ± 36.3 Hz at ambient weather conditions (Stilz & Schnitzler, 2012). If social calls help coordinate movement behaviour, then bats within hearing distance of one another should be influence by the movement and potential recruitment of group members.

Generalized linear mixed effects models were fit in *lme4* with individual as a random intercept nested within social group (Bates, Mächler, Bolker, & Walker, 2015). For models evaluating proportional activity budgets as a response, a binomial family was specified and for all others Gaussian models were used. When a non-linear relationship seemed likely, we fit second-order polynomial models and tested if they fit the data more efficiently than a first order model using the second-order Akaike Information Criterion calculated using *MuMIn::AICc.* The most efficient model was the model with an AICc value at least three units lower than competing models. To evaluate the significance of the fixed effects, we calculated type II p-values using Satterthwaite degrees of freedom method with *lmerTest::anova* for Gaussian models (Kuznetsova, Brockhoff, & Christensen, 2017) and with *car::Anova* for binomial models. To measure the effect size of each Gaussian model, R^2^ was calculated for both the marginal (fixed effects, R^2^_m_) and conditional (fixed and random effects, R^2^_c_) in *MuMIn* (Bartoń, 2016). The full GPS data set and code are available from the Movebank Data Repository (doi released on acceptance, code and data with environmental and behavioural annotation is available at osf.io for review: https://osf.io/d8gaz/?view_only=324382133cfb44c28ada3120b49ea42e).

## Results

### Tracking Summary

The 18 tracked bats mostly foraged in sites that were 20 – 30 km away from their cave roost across the sea (Fig 1), however one of the harem males foraged close to the cave and most individuals showed some indications of quick foraging stops on their return flights to the cave. We found no co-commuting flight within the 41 combinations of tagged pairs of bats. None of the tagged bats moved together to a foraging site in a way that would be consistent with social foraging. While GPS tags had the same programmed on/off time, because of low satellite coverage and late cave emergence, the first GPS record of each tag was 149 ± 49 min after sunset. At this time bats were already commuting and 5.2 ± 6.2 km from the roost when first locations were recorded (range: 350 m – 24.2 km). Each night, bats spent 197 ± 60 minutes outside the roost and travelled 59.2 ± 16.2 km (Table S1). Bats commuted to their foraging areas with slight headwinds at ground speeds of 8.63 ± 2.63 m s^-1^ (airspeed: 9.12 ± 2.69 m s^-1^) and returned to the cave at 7.89 ± 4.08 m s^-1^ (airspeed: 7.76 ± 3.82 m s^-1^) flying with tailwinds, airspeeds that are between their minimum power speed (6.81 ± 0.21 m s^-1^) and maximum range speed (11.0 ± 0.35 m s^-1^). Wind speeds were generally low during the tracking period, with prevailing offshore winds blowing eastward. Bats foraged at ground speeds of 3.95 ± 3.43 m s^-1^ (airspeed: 4.15 ± 3.38 m s^-1^).

**Figure 1.**
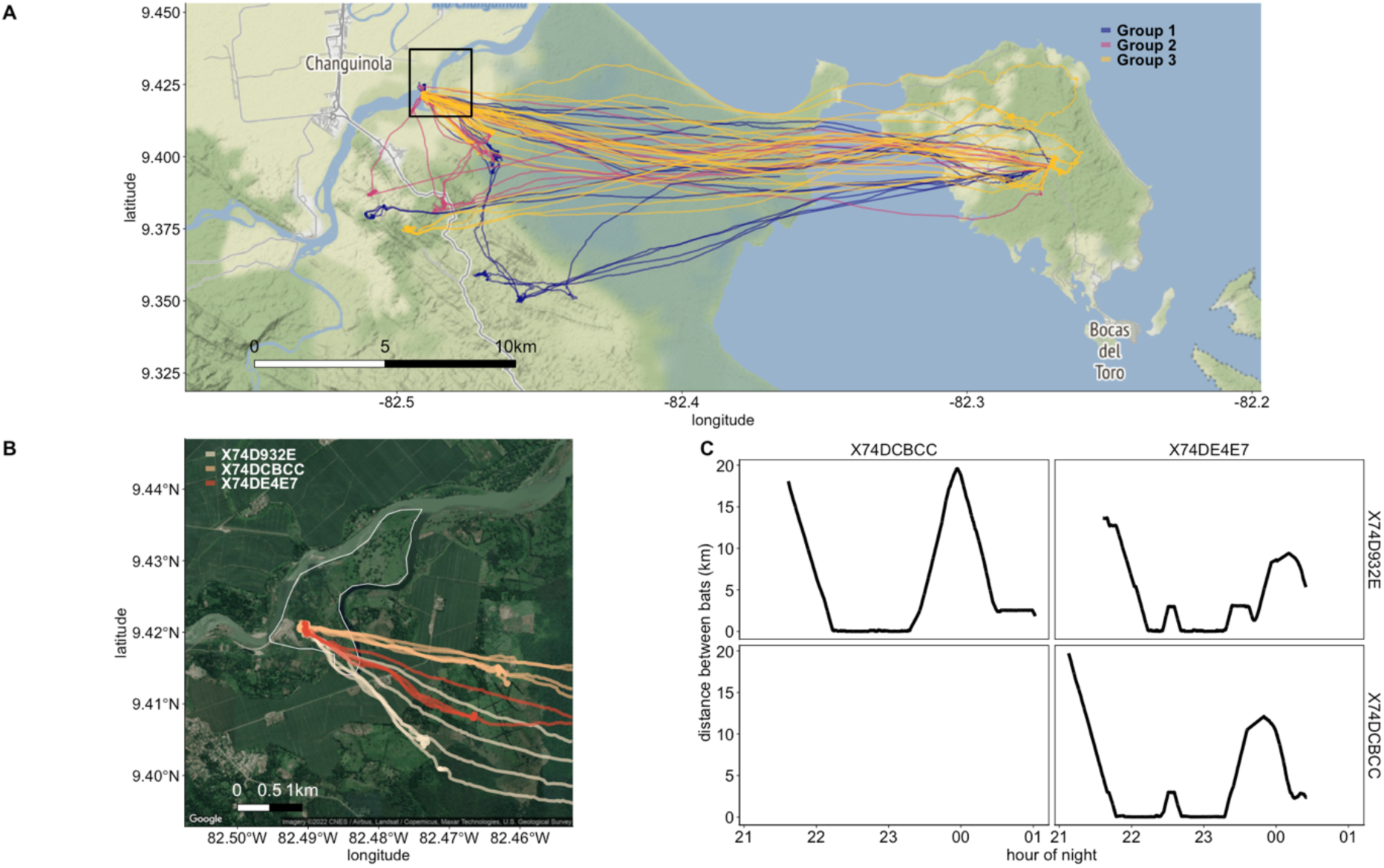
A) Tracking overview with individuals coloured by group membership. B) The island in the Changuinola river (Isla Changuinola) shown with three individuals from group 3 on 2016-03-04, and C) the pairwise distances between these three individuals across that night. Note that in C both axes are repeated to show the simultaneous distance between pairs of individuals and the large distances among individuals at the beginning and end of the nights when commuting. The tracked bats all used similar foraging areas far away from their cave, and no tagged bats commuted together to those sites. Instead, they reunited in the foraging patches 20-30 km from their home cave.

### Behavior & foraging patch use

We ground-truthed the foraging patches on the mainland and found flowering *O. pyramidale* trees in each of them. There were no flowering *O. pyramidale* on Isla Colón, but foraging patches always included flowering *Luehea seemannii*. It is unknown if bats fed on the nectar of *L. seemannii* flowers, or on the animals attracted to this resource. All individuals completed the ca. 25 km commute from the roost to the foraging areas alone (Fig 1C), and individuals were 9.1 ± 5.8 km (mean ± sd) away from one another when commuting. Individuals then converged in the same foraging areas, mostly on the mainland. Individuals used a mix of slow flight, moving and resting during the approximately 200 mins they were outside of the cave, and this did not differ across the three groups (Fig 2A). Bats spent 24.0 ± 19.3% of their time in rest, 24.6 ± 14.6% of time in slow foraging/feeding flights, 31.3 ± 23.6% in faster foraging movements between feeding sites, and 25.1 ± 16.8% of time commuting. To examine social effects on foraging, we only further analysed behaviours other than commuting.

**Figure 2.**
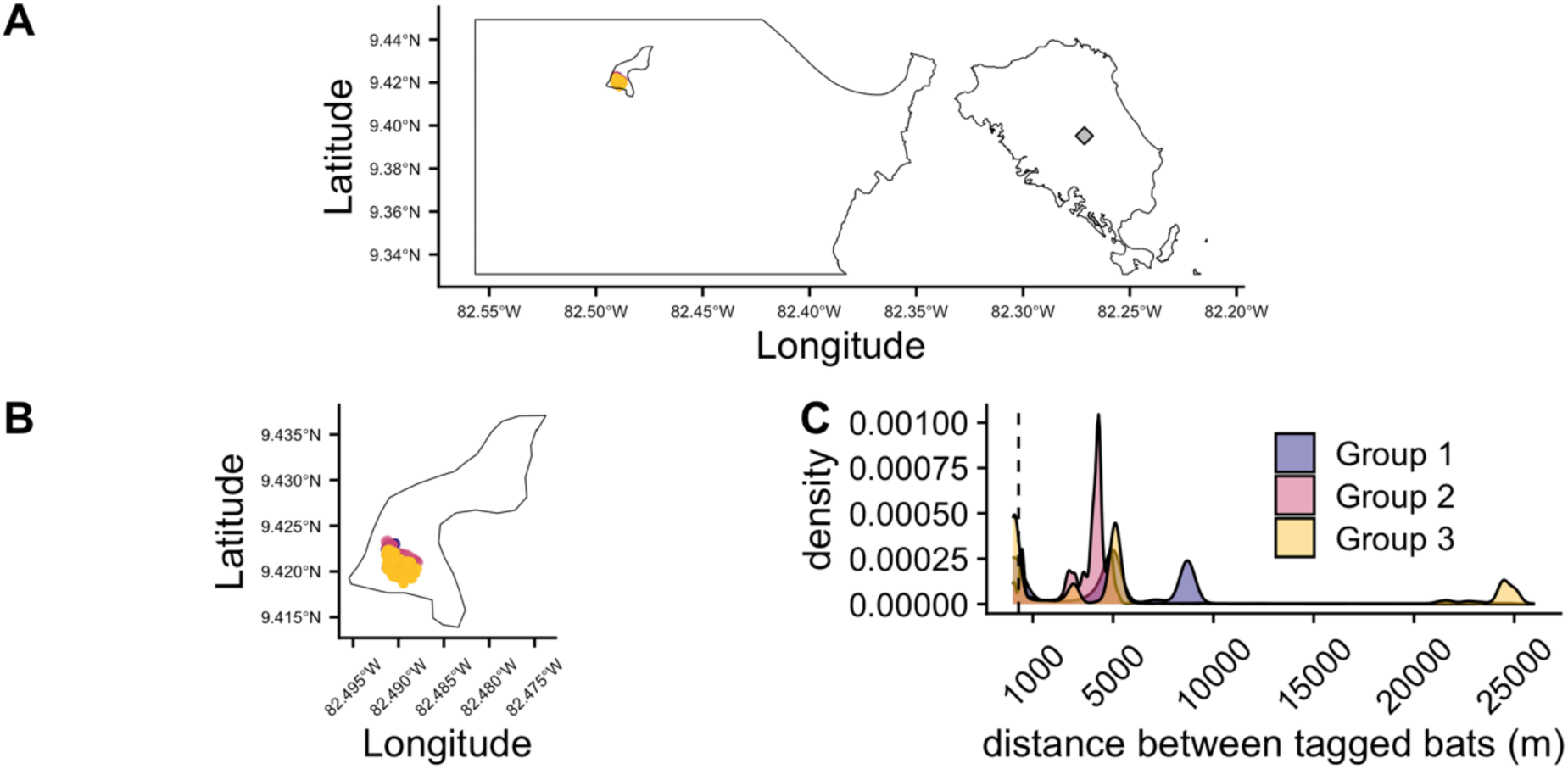
Distances between tagged bats, including the GPS locations that were within the hearing threshold of another tagged group member and assumed threshold for social interaction (290 m) shown for A) the study area, B) Isla Changuinola, and C) total density distributions for all pairwise distances. The capture site is noted with a diamond on Isla Colón (A). All locations within this threshold were found on Isla Changuinola for each of the three study groups. The vertical dashed line in (C) denotes the 290 m transmission threshold of *P. hastatus* social calls.

We identified 73 patches (dense clusters of GPS locoations) that had 100% minimum convex polygon areas of 4 ± 7 ha and likely reflect large aggregations of feeding or roosting trees that were 22.6 ± 31.6 km from the cave roost. Bats used 7.2 ± 4.2 foraging patches (i.e., trees or groups of trees) per night, and patches were 4.0 ± 3.8 km apart. As bats increased their total nightly flight distance, they used more patches (estimate: 0.886 ± 0.034 per additional km, F_1, 20.174_ = 6.861, p = 0.016, R^2^_m_ = 0.13, R^2^_c_ = 0.70; Fig S2). Over the course of the night, bats used 20.72 ± 19.91 feeding locations (flower clusters) (range: 3 - 82). No relationship between the number of feeding locations they visited within the patches and the number of patches used, or with the time they spent outside the roost was detected (Fig S2).

While tagged bats did not commute together to the foraging sites, they would often occupy the same patches as other group members, and proximity to other tagged social group members influenced their behaviour. Bats were within hearing distance of one another (less than 290 m), and presumed threshold of social foraging influence 0.11 ± 0.14% of each night’s tracking (range 0 – 50.0%), and all of these locations occurred on Isla Changuinola for each of the three social groups (Fig 2). Activity budgets differed depending on whether individuals were in the same patch as their neighbours or not (Fig 3A-B, patch estimate F_1, 3_ = 0.028). When in the same patch, individuals were 24.3 ± 21.4 m apart (0.25 – 133 m). The proportion of time spent in rest was larger when the nearest tagged bat was in the same patch as the focal individual, rather than when they occupied different patches that were within 290 m of one another (same patch (median ± MAD): 85.2 ± 18.6%, different patch: 32.2% ± 44.3%, Ξ^2^_3_ = 10.12, p = 0.018, Fig 3A). Bats varied in how much they synchronized behaviours with a nearest neighbour (Ξ^2^_1_ = 4.08, p = 0.043, Fig 3B). Nearest neighbours within a patch were more likely to synchronize resting than other behaviours (Fig 3B). Bats rested 19.3 ± 14.3 m away from their nearest tagged neighbour (range: 0.47 m – 25 km), and resting bouts were longer with decreasing distance from one another, especially within three meters or less (power curve / Freundlich equation intercept = 2.95 ± 0.24 m SE, p < 0.001, Fig 4).

**Figure 3.**
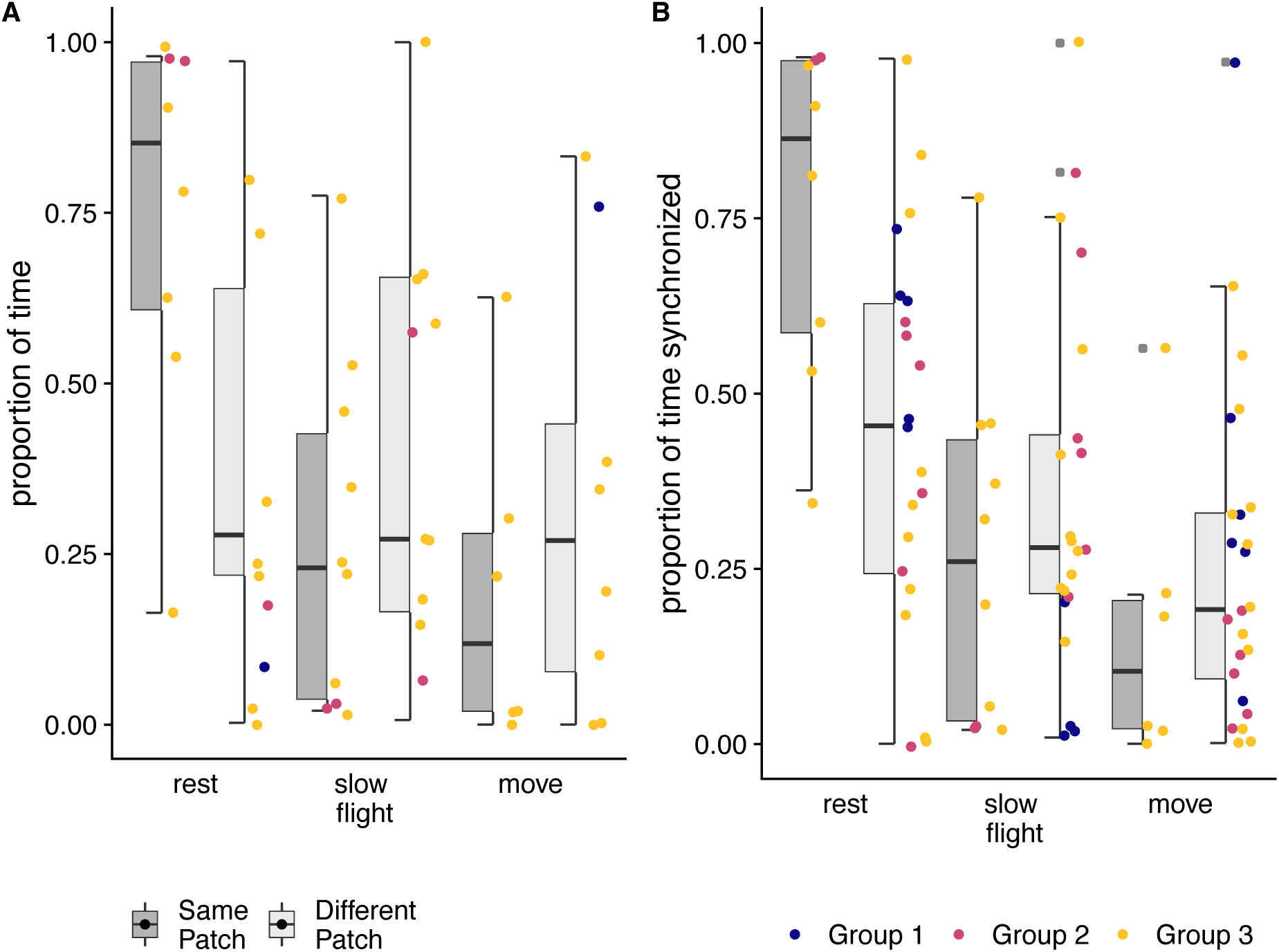
Bat nightly activity budgets and the synchronization of behaviour differ when nearest tagged neighbours are in the same patch or a different patch. A) Bat activity budgets when nearest neighbours occupied the same and different patches, but were within 290 m of one another. B) Behavioral synchrony of nearest neighbours in and out of the same patch. Box plots show proportion of time engaged in behaviors for each patch category, and points show the proportions for each individual bat per night. Tagged bats tended to rest more and at the same time as their nearest tagged individual when they occupied the same patch.

**Figure 4.**
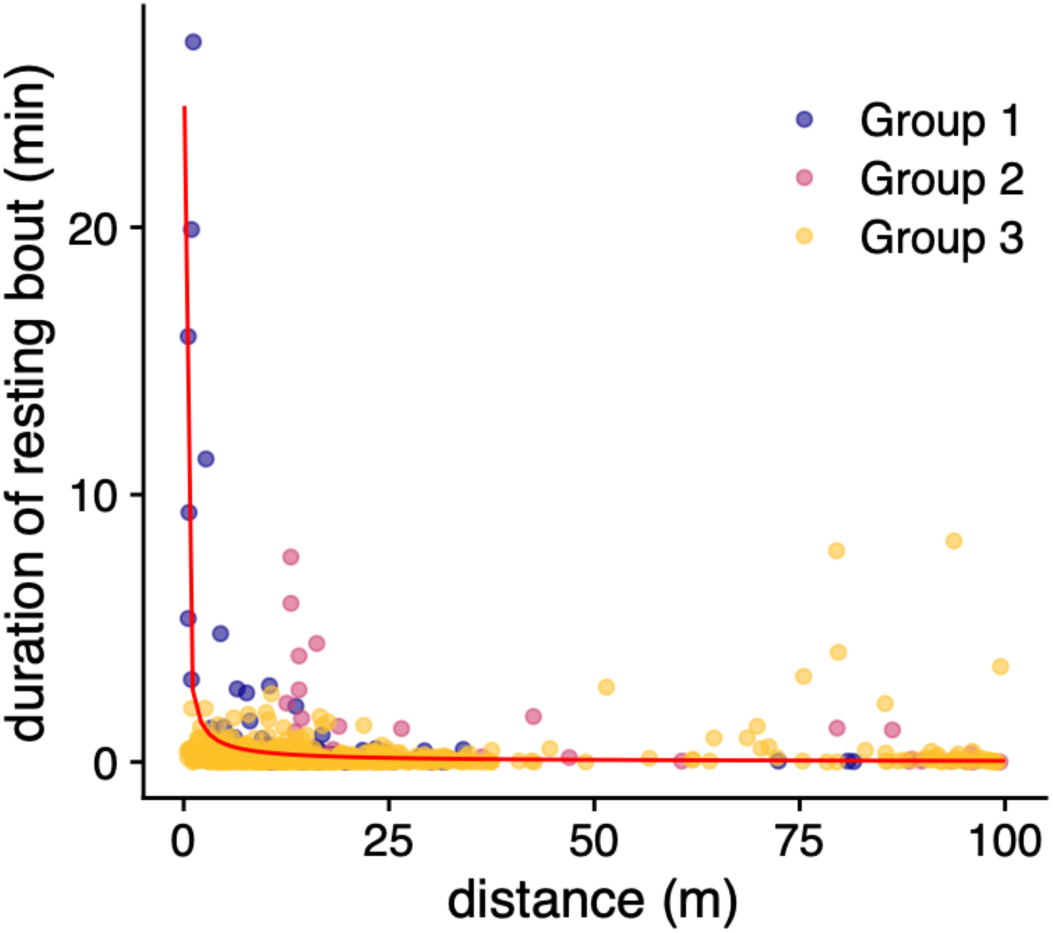
Duration of continuous resting bouts relative to the distance to a bat’s nearest neighbour.

Despite groups being tracked on different nights, the same resting areas tended to be used by bats across all social groups, regardless of the tracking night. We identified 43 resting areas, and one of these resting locations on Isla Changuinola was used by 11 bats over five different nights (Fig S3). Three other locations on Isla Colón were used repeatedly by a single bat over two nights. The remaining 39 sites were used by one bat on one night each.

### Energy costs & feeding requirements

Estimated daily energy expenditure (DEE) was 198.47 ± 69.44 kJ day^-1^ and increased with tracking time (estimate: 0.729 ± 0.174, F_1,25.61_ = 17.462, p < 0.001, R^2^_m_ = 0.383, R^2^_c_ = 0.448, Fig 5A). This is similar to DEE estimates based on allometric estimates from body mass alone of 153.02 ± 5.96 kJ day^-1^ (range: 143.07 – 164.89 (Speakman, 2005)), and to DEE derived from doubly-labelled water measurements in *P. hastatus* that were 36% smaller than those at our study site (76.8 ± 6.6 g vs 121.5 ± 7.6 g in this study, estimates from Kunz et al (1998) are shown in blue in Fig 5A).

**Figure 5.**
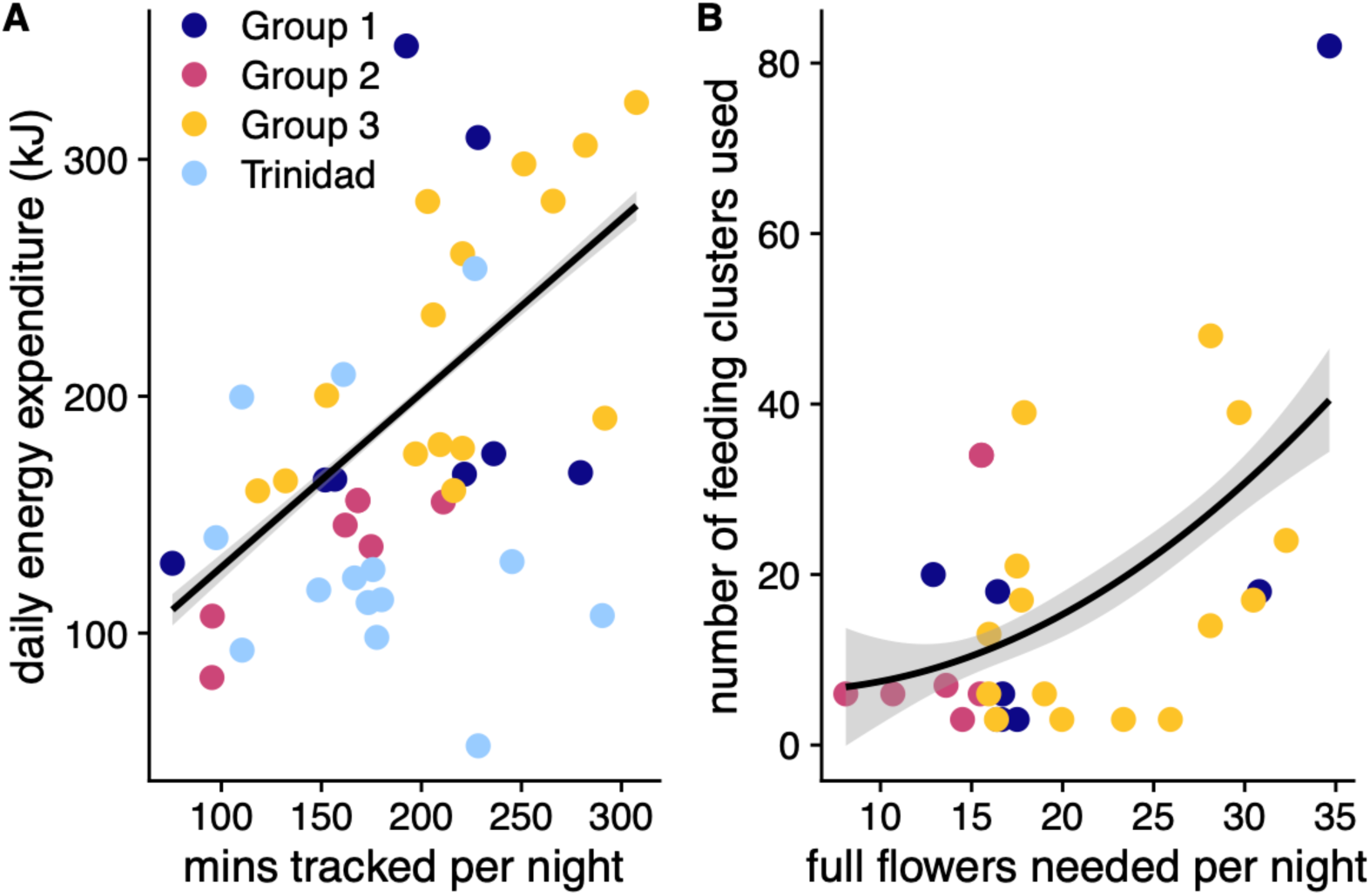
Estimated daily energy expenditure and the estimated energetic returns from foraging. A) Daily energy expenditure and the minutes tracked per night, with comparative data from smaller *P. hastatus* in Trinidad shown in blue (Kunz et al., 1998). The line shows the relationship derived from the individuals tracked in this study. B) The number of flowers needed, based on average energy content, to support the energy requirements of an individual’s estimated energy expenditure predict the number of feeding clusters used per night.

The proportion of time spent less than 290 m from other individuals did not predict total daily energy expenditure (Ξ^2^_1_ = 2.07, p = 0.15) or the number of feeding clusters visited by individuals (Ξ^2^_1_ = 0.30, p = 0.58). We converted the estimated energetic needs for each individual into the number of full *O. pyramidale* flowers required to support them we found that individuals fed from an increasing number of flower clusters as their total estimated energy needs increased (second order polynomial estimate = 55.45 ± 14.89, 27.73 ±14.86, F_2,26.45_ = 8.84, p = 0.001, R^2^_m_ = 0.376, R^2^_c_ = 0.494, Fig 5B). Estimated energy expenditure, as measured through movement, more strongly depended on the total time tracked per night than on the proximity relationships among individuals in a social group, and these closer foraging distances did not predict foraging efficiency.

## Discussion

Based on previous work (Wilkinson & Boughman, 1998), we predicted that *Phyllostomus hastatus* would commute to foraging patches together and forage with their social group to feed on *O. pyramidale* flowers. Instead, we did not find any tagged individuals commuting together to the foraging patches. They commuted over long distances that included the ocean and large commercial banana plantations – landscapes that have few available resources to these bats. Tracked individuals then used the same foraging patches, but on different feeding locations within the patch. When individuals were near group members, they tended to rest, and resting bout duration increased with closer proximity to others.

Closer distance between individuals did not decrease daily energy expenditure or increase foraging success. It appears that while a driving benefit of social foraging is often assumed to be increased foraging success (Giraldeau & Beauchamp, 1999; McInnes et al., 2017; Snijders et al., 2021), this did not seem to be the case under the resource conditions during our tracking study. Instead, resting in *P. hastatus* groups may reinforce social bonds or has benefits outside of foraging efficiency such as predator detection.

Social foraging should increase foraging efficiency, either because food patches are detected more efficiently or because of social facilitation increasing feeding rates. With increasing group size, Trinidad guppies decrease the time it takes to locate food patches and increase their intake rates (Snijders et al., 2021). Larger guppy groups act as more efficient sensors, but this comes at a cost of perceived feeding competition that drives increased bite rates. Individuals of many bat species forage socially to eavesdrop on feeding calls when resources are ephemeral and searching costs are high (Egert-Berg et al., 2018; Fenton, 2003), and some species show extraordinarily coordinated group foraging for these ephemeral resources (Dechmann et al., 2009; Dechmann et al., 2010; Kohles et al., 2022). We found no effect of proximity between group members on time spent foraging, the number of foraging patches or the number of feeding locations.

Astonishingly, we found that individual *P. hastatus* from multiple social groups used the same patches of *O. pyramidale*, but that they commuted over 25 km from their roost without other tagged group members. *Phyllostomus hastatus* in Trinidad were previously found to forage within 10 km of their roost (McCracken & Bradbury, 1981; Williams & Williams, 1970). Displacement studies showed that most *P. hastatus* individuals would successfully navigate back to their home roost after displacement of up to 20 km. At 30 km or more, bats often failed completely to return home (Williams & Williams, 1970). This indicates that the distance at which our study animals were foraging from their cave was at the edge or even outside the range they were familiar with. They had to cross the sea and large commercial banana farms to reach the *O. pyramidale* foraging locations on the mainland. During our resource ground-truthing we found that on their home island and elsewhere south and east on the mainland *O. pyramidale* was not yet available. This suggests that once an isolated *O. pyramidale* patch is discovered, especially early in the flowering season, social information about this quickly moves through a colony. Once they have located it, group members commute to the patch alone. *Phyllostomus hastatus* feeds on various food sources throughout most of the year, but almost exclusively switches to *O. pyramidale* when it is available during the dry season (Wilkinson & Boughman, 1998). It is unknown if this is due to a shortage of other food sources, or to a distinct preference for nectar. However, locating this widely distributed food source may rely on social information transfer (Kohles et al., 2022). This may be a rare, but crucial resource during the transition period when few trees are flowering, and finding food is more unpredictable. Further tracking, mapping of the resource landscape, and detailed dietary analysis are needed to identify how and when bats switch between food resources and the behavioural and energetic correlates of this change.

While they commuted without other tagged individuals, we found that some bats would reunite with social group members in these patches, while others tended to forage completely on their own. This is similar to vampire bats (*Desmodus rotundus*) that fly individually to cattle approximately 300 m away where they then feed with partners with whom they have close social relationships (Ripperger & Carter, 2021). Previous radio tracking of *P. hastatus* in Trinidad found that during most of the year, social group members foraged alone and did not depart or arrive at the cave together, but on rare occasions they joined a group mate in a nearby foraging area (McCracken & Bradbury, 1981). Members of social groups occupied adjacent foraging ranges and social groups were segregated across the landscape (McCracken & Bradbury, 1981), unlike the large overlap we found during this study. Groups in Trinidad were more likely to depart together, and females from the same social group were captured around a *O. pyramidale* feeding site more often than randomly expected during the dry season (Wilkinson & Boughman, 1998). There also appears to be strong social attraction among Trinidad *P. hastatus* groups. Social calls broadcast at flowering *O. pyramidale* attracted bats, and social calls from flying *P. hastatus* were most often noted 3-4 hours after sunset during foraging (Wilkinson & Boughman, 1998). The higher spatial and temporal resolution of our GPS tracks now indicates they are at least not only attracted to the possibility of food resources, but that they may form resting associations while outside of their roost. These resting areas appeared to be conserved across our sequentially-tracked social groups and could indicate that bats from the caves on Isla Colón tend to rest in larger aggregations during foraging. This may keep groups together, but also could have strong anti-predator benefits. Further work targeting these foraging and resting sites through acoustics or thermal tracking would give further insight into the behaviour of these groups away from their roosts.

There could be regional or population differences in the main drivers on social group formations depending on the resource landscape. In Trinidad it has been hypothesized that recruitment of group members to flowering trees may predominantly help bats defend trees against competitors (Wilkinson & Boughman, 1998). In Panamá, however, large animals that bats cannot defend against, such as kinkajous and opossums are the main visitors of *O. pyramidale* flowers (Kays et al., 2012). A flowering *O. pyramidale* with mean peak flower availability of 60 flowers per night provides approximately 600 kJ of energy at the beginning of the night. This is before nectar pools are depleted and trees begin producing less energy-dense nectar (Kays et al., 2012). We estimated that our tracked bats expended 198 ± 69 kJ / day, indicating that a single *O. pyramidale* crown could support the daily energy needs of only 3 - 7 bats. Such a limited resource may be worth defending from conspecifics if all available flowers could be fully exploited (Wilkinson & Boughman, 1998), but a single tree would not support the needs of an entire social group of bats and large clusters of trees would be needed to supply a social group’s daily energy needs. This suggests that a collective resource defence is not the likely explanation for foraging near group mates for this population in Panamá.

While foraging success during this tracking study did not appear to rely on social foraging, there are other highly relevant reasons for individuals in a social group to associate with one another. Within social groups that are structured at least partly by kin relationship, strong social bonds between related individuals have numerous life history advantages (Silk, 2007), extending lifespans (Barocas, Ilany, Koren, Kam, & Geffen, 2011; Silk et al., 2010), success in group conflicts (Samuni, Crockford, & Wittig, 2021), and individual reproductive success (Frere et al., 2010; Schülke, Bhagavatula, Vigilant, & Ostner, 2010; Silk, Alberts, & Altmann, 2003). On Trinidad, however, female *P. hastatus* form stable, relatively closed groups of unrelated females that stay together for their lifetime and show highly developed social bonds (McCracken & Bradbury, 1981; Wilkinson et al., 2016). They develop group-specific social calls (Boughman, 1997; Wilkinson & Boughman, 1998), and recognize and guard group-members’ offspring (Bohn et al., 2009; Bohn, Wilkinson, & Moss, 2007). Low reproductive rates and high infant mortality in this species (Stern & Kunz, 1998) are strong selective pressures on potential cooperation among unrelated females. The ecological and physiological conditions structuring *P. hastatus* social groups may be similar to groups of unrelated females in wild equids (Cameron, Setsaas, & Linklater, 2009) and some cooperative breeding birds (Riehl & Strong, 2018). The exact purpose of this close resting behaviour while foraging, and whether this is a common phenomenon or a result of exceptional circumstances in this season and population, remain unknown and warrants further study.

Social information may mainly be used for the discovery of feeding areas. Commuting to repeatedly used foraging patches 25 km or more from a central roost may not be unusual for some bats (Calderón-Capote et al., 2020; Goldshtein et al., 2020; Harten, Katz, Goldshtein, Handel, & Yovel, 2020; O’Mara et al., 2021), but still presents navigational and energetic risks due to the long distances travelled and potential weather hazards. Nectar and fruit-feeding fruit bats appear to have large and robust cognitive maps of their foraging ranges (Harten et al., 2020; Toledo et al., 2020). These species will all commute from a common roost to distant foraging patches, but there is little to no overlap among individual foraging ranges in animals from these roosting aggregations, and these non-overlapping ranges may be developed through reinforcement learning that minimizes competition (Goldshtein et al., 2020).

Further work mapping resources as seasons change and social groups reach decision points to alter movements will help us further elucidate the intricate relationships between social and foraging behaviour and its energetic context. This may have strong variation among populations that differ in the resource landscapes they encounter but may allow us to better understand the links between species ecological niche and how sociality responds to resource environments and the need for information use.

## Acknowledgements

We would like to thank the owners of Cueva La Gruta for access to the research site. Maurice Thomas, and the Smithsonian Tropical Research Institute assisted with logistics and support, particularly Rachel Page, Rachel Collin, Plinio Gondola, and Urania González. We would also like to thank Gary McCracken, Camila Calderón, Jenna Kohles, Anne Scharf, Mariëlle van Toor, and Kamran Safi for suggestions and insight, and Martin Wikelski and the Max Planck Institute of Animal Behavior (formerly Ornithology) for exceptional support. This work was approved by the Ministerio del Ambiente, Panamá (SE/A-96-15), and the Animal Care and Use Committee at the Smithsonian Tropical Research Institute (2014-0701-2017).

## Supplementary Material

**Table S1.** Biometric and tracking information for the individuals analyzed in this study. Reproductive state is coded as nr: non-reproductive, plac: post-lactation, and nulli: nulliparous. Nightly values for the minutes of tracking and number of locations used are given for means ± sd when more than one night was recorded.

| Bat ID | Group ID | Sex | Reproductive State | Mass (g) | Forearm length (mm) | Wing length (mm) | Wing area (mm<sup>2</sup>) | Nights Analyzed | Time Analyzed (min) | Number of Locations Used | Distance Tracked (km) |
| --- | --- | --- | --- | --- | --- | --- | --- | --- | --- | --- | --- |
| 2016030703 | one | m | nr | 134 | 93.5 | 271.96 | 22966.52 | 2 | 133.99 ± 82.68 | 7296.5 ± 3939.29 | 43.61 ± 35.34 |
| 2016030705 | one | f | plac | 124 | 87.8 | 280.82 | 24543.47 | 1 | 156.67 | 7694 | 42.55 |
| 71A0D95 | one | f | nulli | 122 | 89.9 | 282.74 | 24088.56 | 2 | 224.89 ± 4.82 | 12371.5 ± 1289.06 | 56.38 ± 1.2 |
| 71A111A | one | f | nulli | 109 | 90.8 | 296.86 | 25587.39 | 1 | 279.52 | 16752 | 78.12 |
| 74DDA80 | one | f | nulli | 123 | 91.2 | 278.44 | 23724.58 | 1 | 151.88 | 6268 | 48.52 |
| 74DE9A7 | one | f | plac | 122 | 91.5 | 274.95 | 23752.23 | 1 | 236.13 | 13488 | 65.24 |
| 74D8954 | two | f | plac | 128 | 93.02 | 317.17 | 29422.44 | 1 | 161.88 | 6211 | 53.89 |
| 74DA92F | two | f | plac | 109 | 87.6 | 250.07 | 19667.46 | 2 | 192.87 ± 25.53 | 6767 ± 1236.02 | 70.42 ± 7.41 |
| 74DCA83 | two | f | nulli | 123 | 92.1 | 300.31 | 29703.42 | 2 | 131.85 ± 51.45 | 5028 ± 2224.56 | 56.72 ± 16.26 |
| 74D8C25 | three | f | nulli | 131 | 93.2 | 308.25 | 28074.65 | 1 | 118.15 | 5158 | 37.24 |
| 74D932E | three | f | nulli | 116 | 93.6 | 304.29 | 28397.6 | 2 | 207.63 ± 2.4 | 8783 ± 83.44 | 61.74 ± 6.62 |
| 74DAF9C | three | f | nulli | 126 | 94.5 | 296.29 | 26317.99 | 1 | 95.27 | 5628 | 36.18 |
| 74DCBCC | three | f | plac | 114 | 89.7 | 289.26 | 22936.64 | 2 | 278.75 ± 18.27 | 11910 ± 1578.26 | 67.05 |
| 74DE4E7 | three | f | plac | 124 | 94.2 | 296.11 | 26383.96 | 1 | 197.03 | 6618 | 65.24 |
| 74F7D4C | three | m | nr | 137 | 93.2 | 277.12 | 23586.97 | 1 | 152.63 | 3748 | 27.52 |
| 74F8E19 | three | f | plac | 113 | 91.7 | 289.38 | 25321.02 | 3 | 201.26 ± 61.87 | 7702.67 ± 2597.55 | 62.01 ± 19.62 |
| 74F9F83 | three | f | plac | 116 | 92 | 284.51 | 24720.16 | 3 | 264.17 ± 54.4 | 7866.33 ± 1690.4 | 76.79 ± 10.33 |
| 74FE24E | three | f | plac | 113 | 91.7 | 270.5 | 22057.56 | 2 | 218.35 ± 3.15 | 7528 ± 622.25 | 66.52 ± 2.7 |

**Figure S1.**
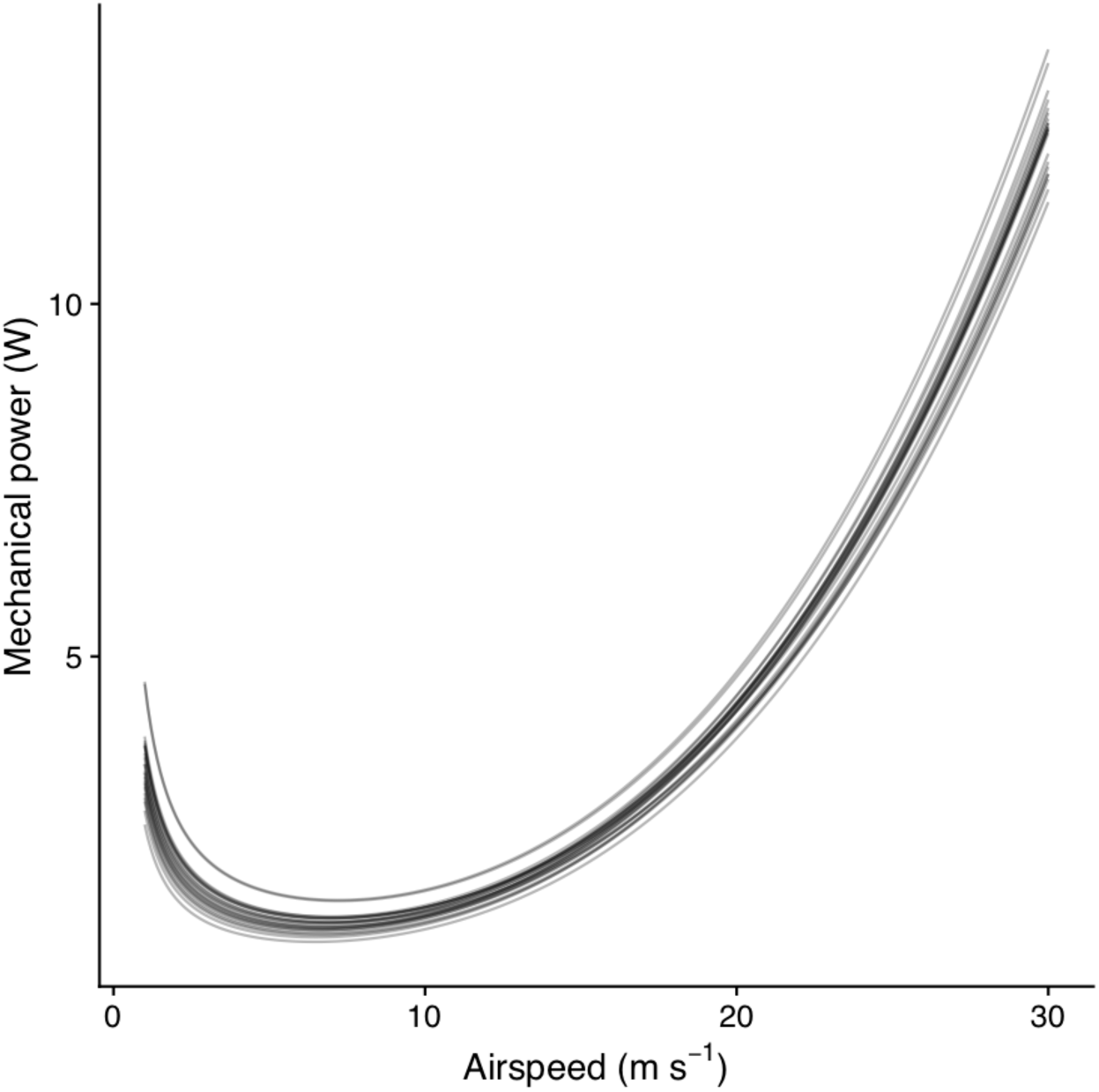
Mechanical power requirements at airspeeds for each bat in this study. Power output was calculated following Pennycuick (2008) at 50 m and 25 °C using each individual’s body mass and wing dimensions.

**Figure S2.**
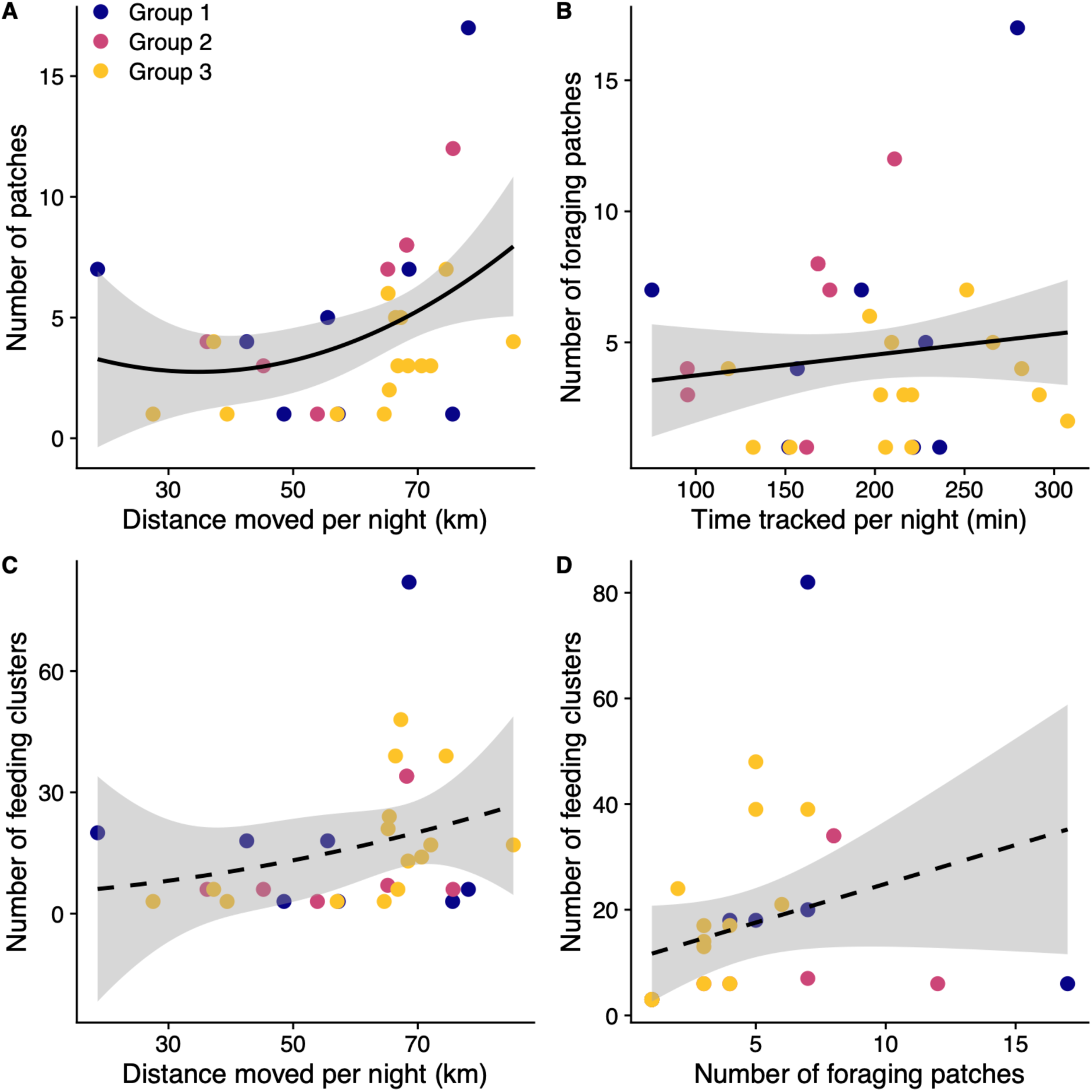
Individual nightly summaries for foraging and feeding. The number of foraging patches used is predicted by A) the distance that individuals moved each night and B) the total time that individuals were tracked each night. The number of feeding clusters identified was not predicted by either C) the distance moved per night or D) the number of foraging patches used. Statistically significant slopes that differed from zero are shown by solid lines, non-significant slopes by dashed lines, and shaded areas indicate 95% confidence intervals.

**Figure S3.**
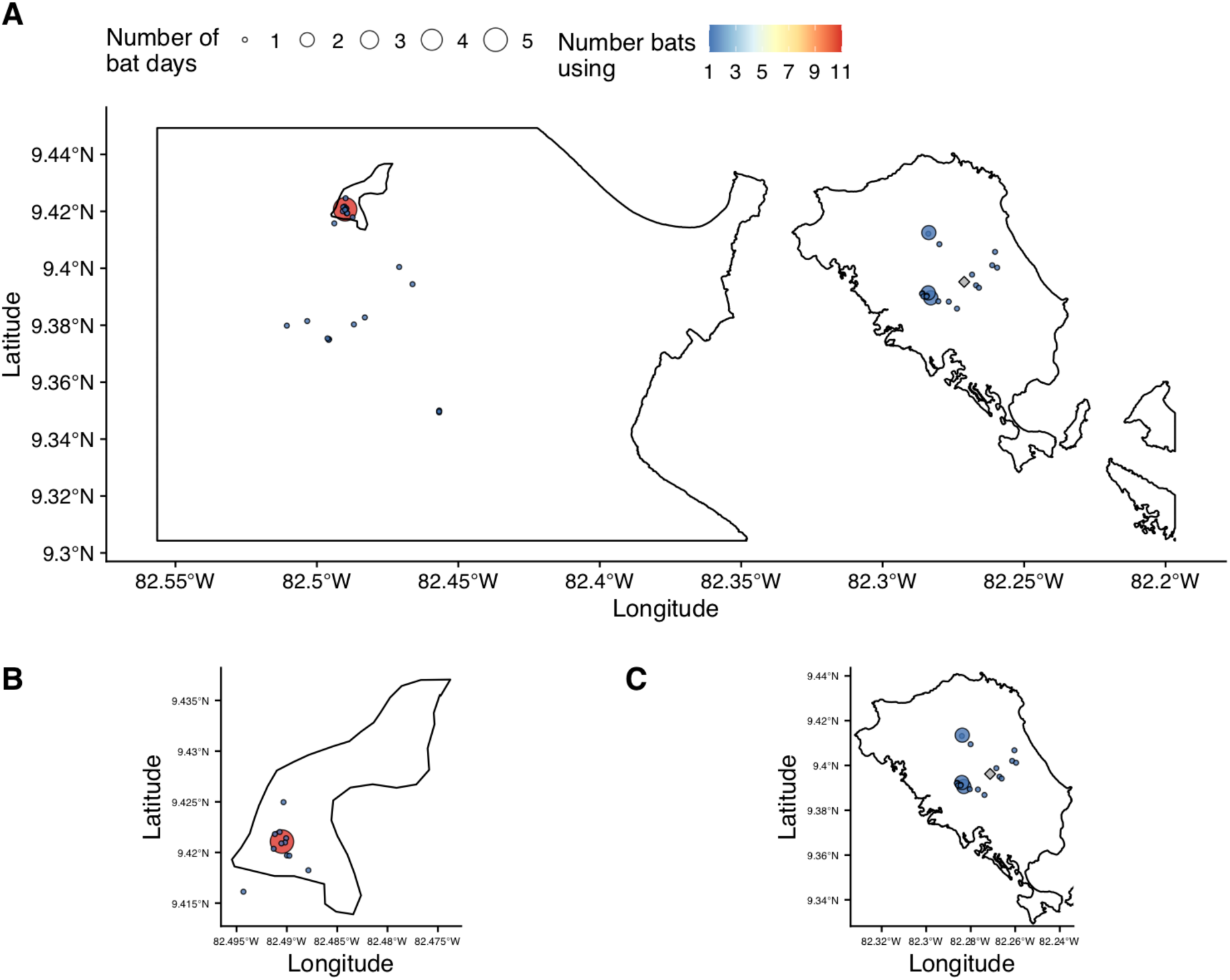
Resting patches clustered from GPS locations. A) Resting patches are shown across the study area, scaled by the number of individual bat days that the sites were used and colored by the number of individual bats that used the patch. Resting patches are shown for B) Isla Changuinola, and C) Isla Colón.

